# The dopamine circuit as a reward-taxis navigation system

**DOI:** 10.1101/2021.04.15.439955

**Authors:** Omer Karin, Uri Alon

## Abstract

Research on certain circuits in simple organisms, such as bacterial chemotaxis, has enabled the formulation of mathematical design principles, leading to ever more precise experimental tests, catalyzing quantitative understanding. It would be important to map these principles to the far more complex case of a vertebrate behavioral circuit. Here, we provide such a mapping for the midbrain dopamine system. Dopamine transmission plays a key role in learning, motivation, and movement, but its systems-level function is not fully understood. We develop a minimal mechanistic model of the dopamine circuit based on physiological and behavioral data, and show that it can be mapped mathematically to the bacterial chemotaxis circuit. Just as chemotaxis robustly climbs attractant gradients, the dopamine circuit performs ‘reward-taxis’ where the attractant is the expected value of reward. The reward-taxis mechanism is based on a circuit feature called fold-change detection, where the circuit outputs the temporal logarithmic derivative of expected reward. The model can explain the general matching law, in which the ratio of responses to concurrent rewards goes as the reward ratio to the power β. It provides an accurate mechanistic value for β as the average gain/baseline ratio of the dopaminergic neurons. Reward-taxis provides testable etiologies for specific dopamine-related disorders.

## Introduction

Foundational work in the field of systems biology showed how, once a biological circuit is well characterized in terms of components and behavior, general mathematical principles can be formed which not only shed light on the original circuit but can generalize to explain many other circuits. It is important to explore whether these principles apply also to a vertebrate behavioral circuit. To do so, it is ideal to begin with a well-characterized circuit in terms of neuronal dynamics, physiology, and behavior. A good candidate for such a circuit is the midbrain dopamine circuit.

Dopamine transmission in the midbrain participates in at least two major biological functions. The first role is the control of movement and motivation (Niv *et al*, 2007; Mazzoni *et al*, 2007; Dudman & Krakauer, 2016; da Silva *et al*, 2018; Shadmehr & Ahmed, 2020). Dopamine transmission regulates movement initiation and vigor on a sub-second level as shown in freely behaving mice (da Silva *et al*, 2018), and defects in dopamine transmission underlie the movement difficulties in Parkinson’s disease (Meder *et al*, 2019).

The second role of dopamine is encoding *reward prediction errors* (Schultz *et al*, 1997). Reward prediction errors are the difference between the experienced and predicted rewards. They play a key role in a specific method of reinforcement learning called temporal difference learning (TD learning) (Schultz *et al*, 1997; Sutton & Barto, 1998; Glimcher, 2011; Steinberg *et al*, 2013). In addition to rapid responses encoding reward prediction errors, dopamine also slowly ramps up when approaching a reward (Howe *et al*, 2013; Hamid *et al*, 2016; Mohebi *et al*, 2019). Importantly for the present study, a recent experiment demonstrated that dopaminergic ramps represent a temporal derivative-like computation on expected reward (Kim *et al*, 2020). Kim et al. used virtual reality to manipulate the rate of change of expected reward as mice moved towards a target, and found that mice responded to the temporal derivative of expected reward, rather than to its absolute value.

To better understand the dopamine circuit on the system level, it is important to unify, in a single conceptual framework, both its reward-derivative and its motion control functions (Berke, 2018; Shadmehr & Ahmed, 2020). Why does dopamine play two important roles at the same time: invigorate movements and encode the derivative of expected reward? Theoretical studies have analyzed this question from the perspectives of learning (Friston, 2010; Bogacz, 2020) and cost-benefit theories (Yoon *et al*, 2018; Shadmehr & Ahmed, 2020). Here, we consider the function of the circuit from the point of view of navigation, a connection anticipated by early work on TD learning (Montague *et al*, 1995).

For this purpose, we develop a mechanistic model of the dopamine circuit, and show that it can be mathematically mapped to the navigational circuit that guides bacterial chemotaxis, a foundational circuit in systems biology. We show that experimental observations on dopaminergic regulation can be explained by a circuit property known as fold-change detection *(FCD)*, in which the output of the circuit responds to *relative* rather than absolute changes in the input (Shoval *et al*, 2010; Adler & Alon, 2018). FCD was discovered in bacterial chemotaxis and was then found to be widespread in sensory and navigational circuits (Adler & Alon, 2018). In the case of dopamine, the input to the circuit is expected reward. Additionally, just as chemotaxis controls bacterial runs and tumbles, we build on measurements of animal movement to propose a model where high dopamine results in prolonged bouts of vigorous and correlated movements (“runs”) whereas low dopamine causes pauses and slow reorientations (“tumbles”). This model for dopaminergic regulation of movement indicates how the two roles of dopamine, reward-derivative and motion-regulator, combine to provide a robust navigation system that climbs reward gradients. We call this the ‘reward-taxis’ model for the dopamine system. The model provides a mechanistic explanation for the well-established generalized matching law of behavior, predicts search dynamics that are invariant to the scale of the rewards, and provides testable etiologies for specific dopamine-related disorders.

## Results

### Dopamine release as fold-change detection of expected reward

We begin by developing a minimal model for midbrain dopamine regulation by environmental cues (Figure 1A). Consider a behaving animal exploring an open field for a reward of magnitude *u*, such as a food or drink. For simplicity, we assume a uniform response amongst all dopaminergic neurons. In reality, there are important heterogeneities between and within midbrain structures, and some dopaminergic neurons may be specialized to specific cues (Eshel *et al*, 2016; Parker *et al*, 2016; da Silva *et al*, 2018; Lee *et al*, 2019; Engelhard *et al*, 2019). If the animal is familiar with the environment, dopamine release *d* will increase as the animal observes cues that indicate closer proximity to the reward (Howe *et al*, 2013; Hamid *et al*, 2016; Kim *et al*, 2020). This is due to the increase in expected reward *R*, which decays with distance from the target (Howe *et al*, 2013; Gershman, 2014; Kim *et al*, 2020). Here expected reward is defined based on TD learning: the expected temporally discounted sum of present and future rewards (see Methods).

**Figure 1.**
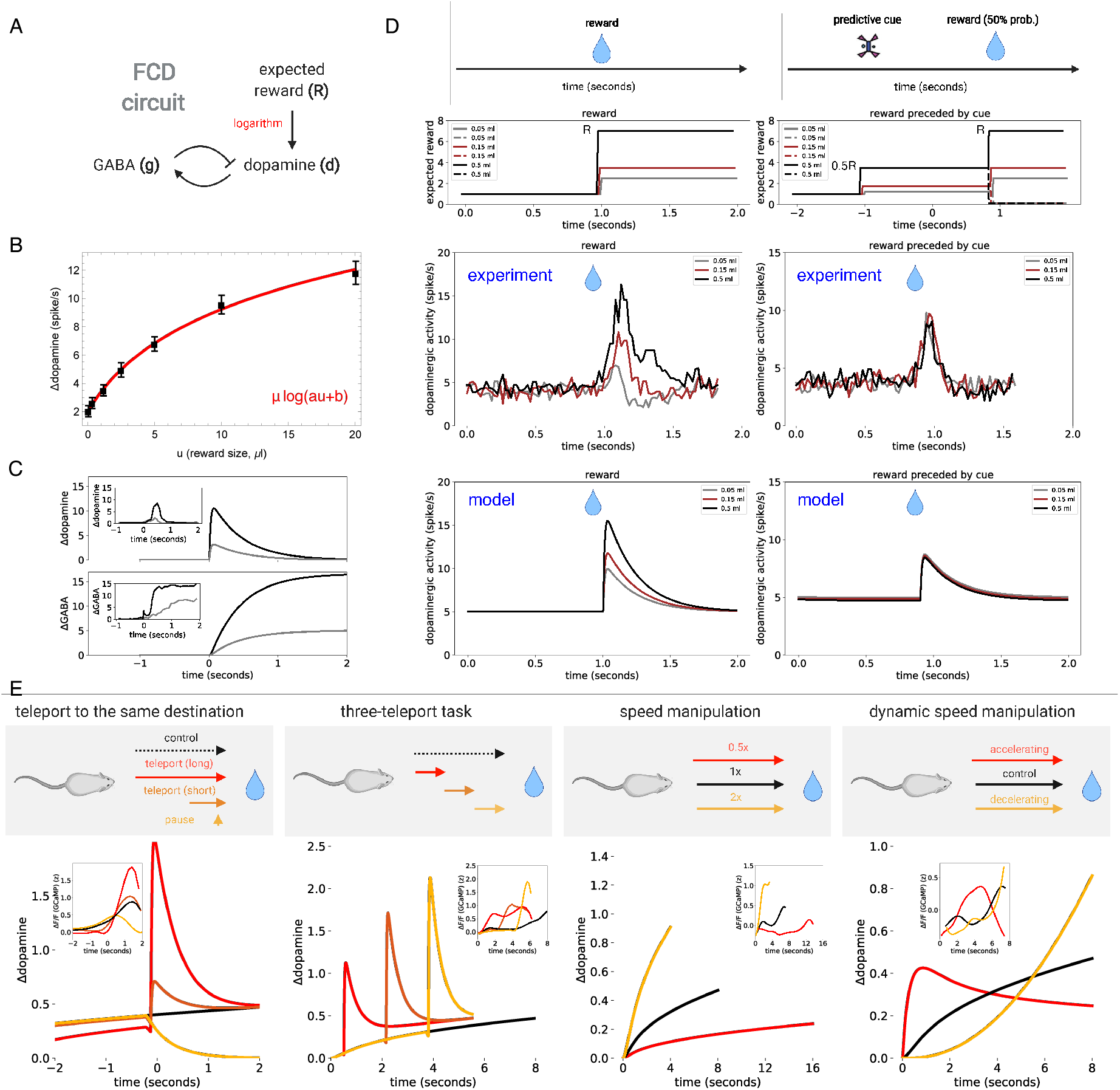
Dopamine responses as fold-change detection of expected reward. (A) Minimal circuit for dopamine responses. Dopamine *(d)* is activated by the logarithm of expected reward (log *R*) and is inhibited by negative feedback from GABAergic input (*g*). See *Methods* for equations and parameters. (B) The response of VTA dopaminergic neurons in mice (n=5) to a water reward of variable volume (black squares, mean ± SEM, taken from Figure 1C in (Eshel *et al*, 2015)) is well-described by the logarithmic relation Δ*d* = *μ* log(*au* + *b*) (*r*^2^ = 0.999, best-fit parameters *a* = 0.5 ± 0.1, *b* = 1.5 ± 0.05, *μ* = 4.9 ± 0.45, N=7 reward magnitudes). (C) Simulation of dopamine and GABA responses to a step increase in expected reward input which corresponds to the presentation of a reward-predicting cue. The step is given by *R*(*t*) = *R*_0_ + *λθ*(*t* – *t*_0_) where *θ*(*t* – *t*_0_) is a unit step function, *R*_0_ = 1 and *λ* = 7 for the black line (large reward) and *λ* = 1.8 for the gray line (small reward). *Insets*. Average change in firing rates from dopaminergic (type I) and GABAergic (type II) VTA neurons, in response to reward-predicting cues for a small reward (gray) or a large reward (black). Data from Figure 2D of (Cohen *et al*, 2012). (D) Population responses of dopaminergic neurons of two Macaque monkeys to variable size liquid reward, either without a preceding cue (left panels, n=55 neurons), or with a preceding visual cue that predicts reward delivery with 50% probability (right panels, n=57 neurons). *Upper*. The expected-reward input following the reward-predicting cue is *R* = 0.5(*b* + *λu*) (where *λ* is proportional to reward magnitude), which is doubled following reward delivery, *R* = *b* + *λu*, where *u* is the reward magnitude (we use *b* = 2 and *λ* = 10*ml*^-1^). *Middle*. Experimentally measured average dopaminergic responses, adapted from Fig. 2A and Fig. 4B of (Tobler *et al*, 2005). When the reward is delivered without a cue, dopaminergic responses increase with reward magnitude (left panel). When it is given after a cue that predicts reward delivery with 50% probability, dopaminergic responses to reward delivery are identical (right panel), as predicted by the FCD property. This is despite the 10-fold difference in reward magnitude. *Lower*. Simulations of the dopamine model capture the experimentally observed dynamics. (E) Dopamine output to movement in a reward gradient given by 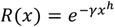, with *h* = 1.5, *x*_0_ = 1, *ν*_0_ = 1 and *γ* = 0.04, and perturbations as described in Figure 2 of (Kim *et al*, 2020). *Insets*. Corresponding dopaminergic outputs from mice (left to right: n=11, n=11, n=15, n=5) VTA neurons measured by calcium imaging, from Figure 2CGKO in (Kim *et al*, 2020), smoothed using a Savitzky–Golay filter. All simulations were performed with the parameters provided in Table 1.

How does dopamine release, *d*, depend on expected reward, *R*? Recordings from VTA dopaminergic neurons in mice reveal a sublinear relationship between the magnitude of received rewards *u* and the dopaminergic response above its baseline, Δ*d*(*u*) (Eshel *et al*, 2015, 2016) (Figure 1B). The sublinear relationship indicates that dopamine neurons may be (at least in some magnitude range) responding to the logarithm of expected reward, i.e., that: Δ*d*(*u*) = *μ* log(*R*(*u*)), where *R*(*u*) = *au + b* is the expected reward, where *a* is a scaling factor and *b* is a magnitude-independent component of the expected reward. The parameter *μ* is the gain of the dopaminergic response. Logarithmic responses are consistent with widespread logarithmic coding in the brain (Dehaene, 2003; Nieder & Miller, 2003; Shen, 2003; Dehaene *et al*, 2008; Nieder & Dehaene, 2009; Laughlin, 1989) as well as with economic notions of utility (Bernoulli, 1968; Rubinstein, 1977).

To test this, we fit the function *μ* log(*au + b*) to the average dopaminergic response to a variable water reward in mice (Eshel *et al*, 2016), finding an excellent fit (*r*^2^ = 0.999), with a gain of *μ* = 4.94 ± 0.45.

In addition to activation by expected reward, dopaminergic neurons in the VTA are inhibited by the output *g* of adjacent GABAergic neurons, and this inhibition is subtractive (Eshel *et al*, 2015, 2016). We therefore propose the following minimal description of dopamine release:

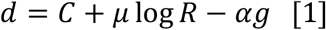

where *C* is the baseline activity of the dopaminergic neurons when log *R* = *g* = 0, *μ* is the gain, *R* is perceived expected reward, and *α* is the effectiveness of inhibition by the GABAergic output *g*. Since our model focuses on the regulation of behavior, rather than on learning or representation, we will assume that the log-transformed expected reward log *R* is an input signal that is provided to the circuit. Additionally, while subtractive inhibition was established for VTA dopaminergic neurons, we assume that similar regulation is shared among all midbrain dopaminergic neurons.

To complete the model requires a minimal description of the dynamics of GABAergic output *g*. The mechanisms of interaction between GABAergic and dopaminergic neurons are complex, and there are many local and remote interactions (Morales & Margolis, 2017; Cox & Witten, 2019). However, there are several experimental observations that impose constraints on these interactions (Figure 1CDE). First, upon presentation of a reward-predicting cue - equivalent to a step increase in *R(t)* - dopamine rapidly increases and then drops and adapts precisely to its baseline on a sub-second timescale (Schultz *et al*, 1997; Tobler *et al*, 2005; Cohen *et al*, 2012) (Figure 1C). Second, dopamine responses are scaled relative to the expected reward baseline: when the expected reward is of magnitude *u* and the received reward is of magnitude *λu* for a constant *λ*, dopamine responses are nearly identical and independent of *u* (Tobler *et al*, 2005) (Figure 1D). Third, dopamine release represents a temporal-derivative-like computation of *R(t)* (Kim *et al*, 2020) (Figure 1E). Finally, GABAergic activity tracks *R(t)* (Cohen *et al*, 2012) (Figure 1C).

These experimental observations are the hallmarks of a circuit feature known in systems biology as *fold-change detection (FCD)* (Shoval *et al*, 2010; Adler & Alon, 2018) (Figure 1A). FCD is a property where the circuit output depends only on relative changes in the input, rather than absolute changes. FCD circuits output the temporal (logarithmic) derivative of low-frequency input signals, whereas high frequency inputs are normalized by their baseline (Tu *et al*, 2008; Adler *et al*, 2014; Lang & Sontag, 2016).

FCD circuits are ubiquitous in biological sensing, as reviewed in (Adler & Alon, 2018). They can be implemented by a handful of specific feed-forward and feedback mechanisms (Adler *et al*, 2017). In all FCD mechanisms, a slow internal variable (*g*) encodes the background input level and acts to normalize the output accordingly. Taken together, the FCD property can explain all of the experimental observations on dopamine responses described in Figure 1. Since we do not know the exact mechanistic implementation of the FCD property in the dopamine circuit, we will make the simple assumption of a negative feedback loop, and choose the simplest design that provides FCD. In this design, inhibitory neuron activity *g* is given by an integral-feedback equation with respect to dopamine release *d*:

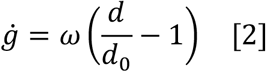

Or, more generally, *ġ* = *F*(*d*) where *F*(*d*) increases with *d* and has a single zero at *d* = *d*_0_. This feedback loop generates exact adaptation of *d*: the only steady state solution is *d* = *d*_0_. *d*_0_ is the homeostatic activity level of dopaminergic neurons, which is about ∼5 spikes/s in mice (Robinson *et al*, 2004; Eshel *et al*, 2016). The parameter *ω* determines the adaptation time of the dopaminergic neurons after a change in *R*(t). This timescale is on the order of hundreds of milliseconds. For the GABAergic neurons, after a step change in *R(t)*, the firing rate in the model increases proportionally to the logarithm of *R(t)*, such that 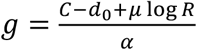 (this is because GABAergic output integrates dopaminergic activity). Taken together, Eqs. 1-2 provide a minimal model for dopamine responses to expected reward inputs *R(t)*.

### A reward-taxis model for dopamine regulation of behavior

To link the dopamine circuit to animal behavior, we provide an additional equation as a minimal model for dopamine control of motion. The equation is motivated by the well-established role of dopamine in the regulation of action vigor and locomotor activity (Niv *et al*, 2007; Beierholm *et al*, 2013; Panigrahi *et al*, 2015; Ek *et al*, 2016; da Silva *et al*, 2018).

We focus on stochastic exploratory motion, as opposed to directed and deliberate motion along a specific path. We therefore model movement as a random-walk process, which is a standard approach for modelling animal movement (Blackwell, 1997; Codling *et al*, 2008; Smouse *et al*, 2010; Mwaffo *et al*, 2015; Hanks *et al*, 2015; Michelot *et al*, 2019). While the model is presented in the context of physical spatial motion of an animal, it can be readily generalized to other aspects of behavior, as we will detail below.

We assume that the navigating animal can be in one of two states – arrested or moving. We also assume that when the animal is moving, it runs in a straight line, and when the animal is arrested, it chooses its subsequent movement direction at random (allowing the new direction to be correlated with the previous direction does not affect the conclusions). Such a model can capture, for example, the swimming behavior of zebrafish (Budick & O’Malley, 2000; Mwaffo *et al*, 2015), or the movement of mice in an open field (da Silva *et al*, 2018). The random walk has two parameters: movement speed *v* (units of length/time), and run duration, which is exponentially distributed with mean *ϕ* (units of time).

Dopamine increases movement vigor, which may be due to increasing *v* or increasing *ϕ*. In zebrafish, the dopamine agonist apomorphine increased the speed of swimming bouts at low doses (higher *v*), and the duration of swimming bouts (higher *ϕ*) at high doses (Ek *et al*, 2016). In a recent study, da Silva et al. performed sub-second measurements of SNc dopaminergic neurons in freely behaving mice in an open field (da Silva *et al*, 2018). The neuronal measurements were coupled with high-resolution motion measurements. Dopamine activity prior to movement increased the probability and vigor of subsequent movement (da Silva *et al*, 2018). Both these effects may again manifest in higher movement speed *ν*, or in correlated movement patterns that correspond to higher *ϕ*.

Based on these observations, we propose two possibilities for dopamine regulation of movement. In the first possibility, dopamine increases run duration:

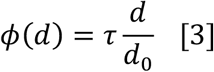

Where *τ* is the average run duration in an environment without reward cues. Dopamine *d* in Eq. 2 corresponds to the (normalized) dopaminergic activity in the brain, so loss of dopaminergic neurons (as occurs in Parkinson’s disease) reduces *d* proportionately, which effectively reduces run duration *τ*. We assume a linear relation between run duration and *d*, which is consistent with the gradual increase in movement vigor with dopaminergic activity (see Fig. 1j-k in da Silva et al. (da Silva *et al*, 2018)). The important aspect is that high dopamine results in directed motion, whereas low dopamine decorrelates motion.

We can also consider an alternative model where dopamine increases running speed:

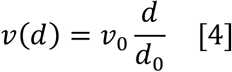

In this case, loss of dopamine neurons effectively decreases the average running speed *ν*_0_. Such a speed-control model provides the same conclusions as a run-duration-control model with Eq. 3 (Methods). However, in order to emphasize the analogy with chemotaxis, we use Eq. 3. Note that in both cases, we focus on the sub-second regulation of movement parameters by dopamine, as observed by da Silva et al. (da Silva *et al*, 2018).

Eqs. 1-3 define a biased random walk that is guided by the gradient of the logarithm of the expected reward (Figure 2B), which we call *reward-taxis*.

**Figure 2.**
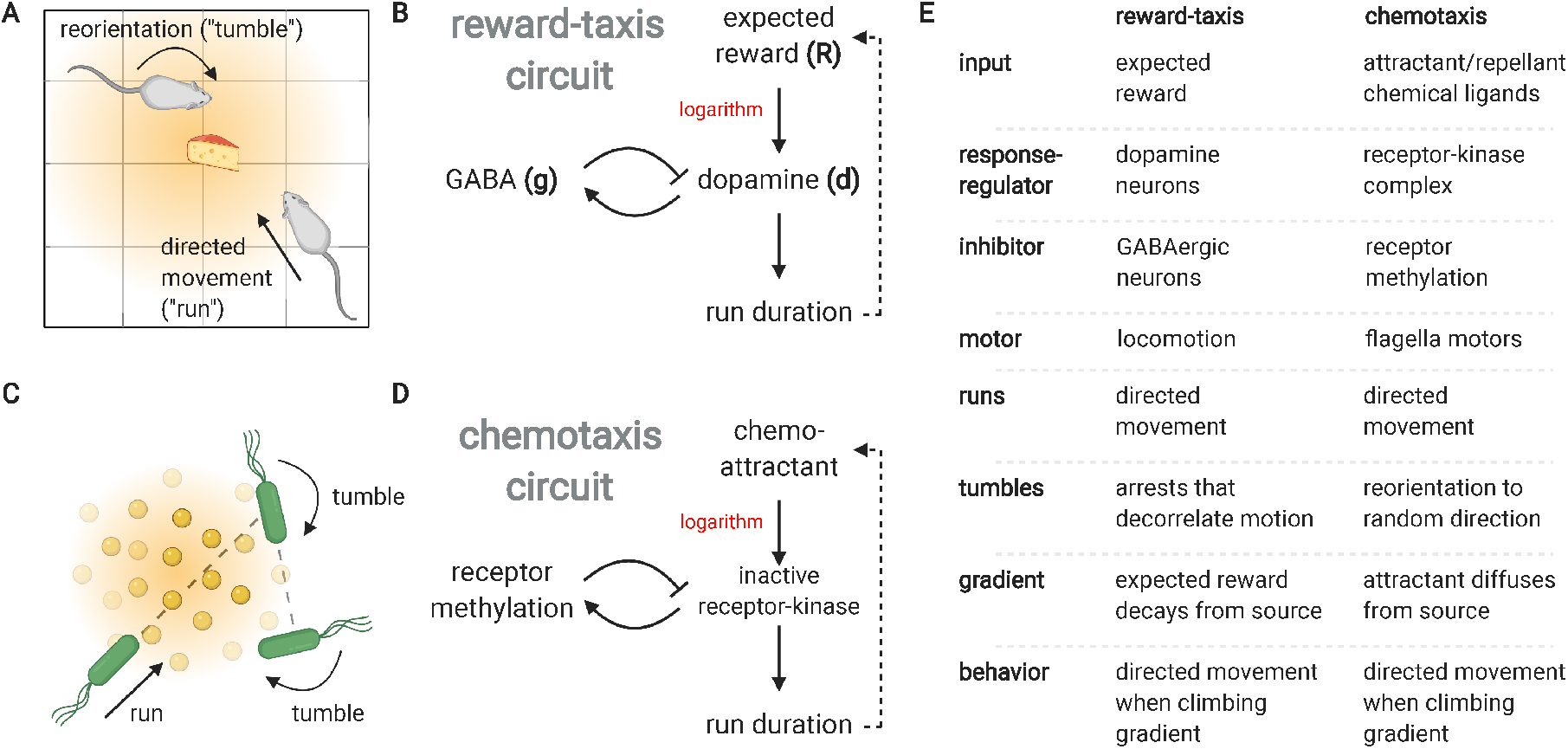
Dopamine regulation of behavior in the model is analogous to bacterial chemotaxis. (A) The behavior of an animal in an open field is modelled as a series of directed movements (runs). The direction of each run is chosen at random (or, more generally, stops between runs decorrelate motion direction), and the duration of each run increases with dopamine level. (B) Dopamine is controlled by an FCD circuit activated by expected reward. (CD) The reward-dopamine-behavior circuit is analogous to the chemotaxis circuit that underlies bacterial navigation towards chemo-attractants. Bacterial motion is composed of a series of runs. The direction of each run is randomized by tumbling events, and run duration increases with the inactivation of a receptor-kinase complex, which is controlled by an FCD circuit activated by chemoattractant concentration. (E) Table detailing the mapping between the dopamine system and the chemotaxis system.

**Figure 3.**
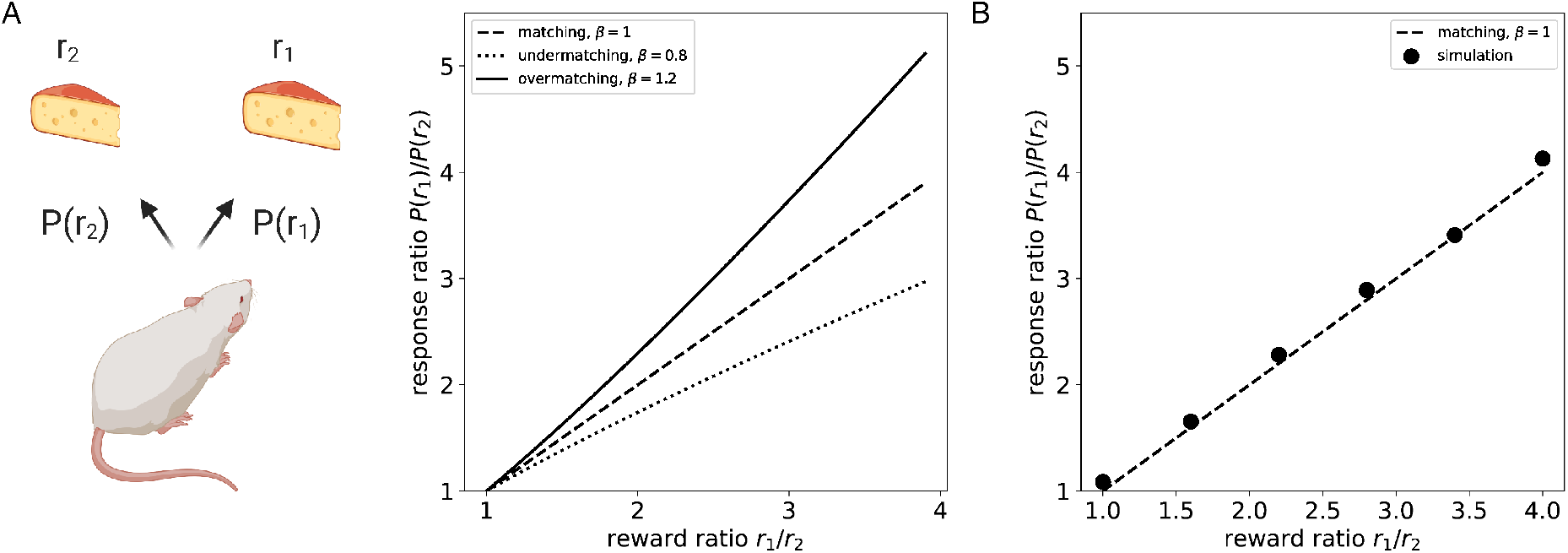
Reward-taxis model provides the generalized matching law. (A) The generalized matching law, amply supported experimentally, is a power-law relation between the frequency of rewards and the frequency of responses 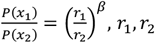, are rewards at locations *x*_1_, *x*_2_. Perfect matching occurs for *β* = 1, while undermatching/overmatching are due to variations in *β*. Such a power-law relation arises directly from the dynamics of the model where 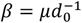. (B) Simulation of the dopamine model for choice between two rewards *r*_1_, *r*_2_. Expected reward is the sum of two Gaussians: 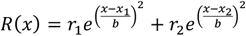 (the exact distribution is not important for matching) with *τ* = 100*ms*, μ = 5, *d*_0_ = 5, ν = 25*cm s*^−1^, *x*_1_ = +30*cm, x*_2_ = −30*cm, b* = 20*cm* (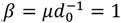, Methods). The model was simulated for different ratios of *r*_1_/*r*_2_, and the response ratio was estimated by the ratio of the time spent ±2.5*cm* from *x*_1_, *x*_2_.

This biased random walk is analogous to bacterial chemotaxis: bacteria such as *E. coli* use a run-and-tumble navigation strategy to climb gradients of chemical attractants (Figure 2CDE) (Berg & Brown, 1972; Tu *et al*, 2008; Shoval *et al*, 2010; Sourjik & Wingreen, 2012). The chemotaxis signaling network is based on an FCD circuit that controls run duration (Shoval *et al*, 2010; Lazova *et al*, 2011). It therefore maps onto dopamine release dynamics, where expected-reward inputs play the role of chemoattractant concentration in chemotaxis. Both systems respond to relative input levels. They both respond to the derivative of low-frequency input components and respond proportionally to high-frequency input components (Tu *et al*, 2008), and show exact adaptation to step changes in input. Thus, the dopamine circuit can allow animals to climb gradients of reward expectation.

The reward-taxis model was presented for whole-body spatial movement, but its assumptions are general and may potentially extend to other aspects of behavior. One such aspect is *hippocampal replay*, the activation of hippocampal place cells during sharp-wave ripples (Gomperts *et al*, 2015; Ólafsdóttir *et al*, 2018; Stella *et al*, 2019). Hippocampal replay consists of a firing sequence of neurons that represents temporally compressed motion trajectories (Lee & Wilson, 2002; Pfeiffer & Foster, 2013; Ólafsdóttir *et al*, 2018). It can occur either during sleep or rest (“offline”) or when the animal is awake and engaged in a task (“online”). Online replay plays an important role in planning and navigation (Pfeiffer & Foster, 2013). The activation of the place neurons corresponds to stochastic movement trajectories with a characteristic speed (Davidson *et al*, 2009) that are biased towards the location of rewards (Pfeiffer & Foster, 2013); when foraging is random, the trajectories are diffusive, resembling Brownian motion (Stella *et al*, 2019). Hippocampal activity during online replay is tightly coordinated with reward-associated dopamine transmission (Gomperts *et al*, 2015). To map reward taxis to hippocampal replay requires that dopamine transmission modulate the stochastic trajectories of hippocampal replay, for example through the modulation of reorientation frequency.

### Reward-taxis quantitatively provides the matching law of operant behavior

We next ask how the reward-taxis model compares with features of animal behavior. We find that the model provides the *general matching law of operant behavior*, one of the best-established behavioral phenomena (Herrnstein, 1970; Baum, 1974, 1981; Baum *et al*, 1999; Lau & Glimcher, 2005; McDowell, 2013). If an animal has a choice of two rewards with expected magnitude *r*_1_, *r*_2_ at two locations *x*_1_, *x*_2_, the long-term average of its response rates (the relative amount of time it chooses each reward (Baum & Rachlin, 1969)) *P*(*x*_1_), *P*(*x*_2_) goes as a power *β* of the ratio of rewards:

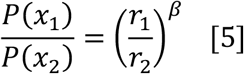

The matching law holds both for varying reward magnitude and varying reward probability (Davison & McCarthy, 1988). The matching law was originally proposed with perfect matching *β* = 1 (Herrnstein, 1961). A large number of studies in various vertebrate species under different experimental conditions observed that *β* can be somewhat variable, showing slight undermatching (*β* < 1) and overmatching (*β* > 1), with the former more commonly observed (Baum, 1974; William, 1979; Baum, 1981; Davison, 1991; Baum *et al*, 1999; Lau & Glimcher, 2005; McDowell, 2013). Matching is a robust property which holds over orders of magnitude of reward ratios (up to ∼1:500 in pigeons (Baum *et al*, 1999)). The overall robustness of the matching law has led authors to suggest that it reflects intrinsic properties of the vertebrate nervous system (McDowell, 2013).

While previous explanations for the general matching law focused on underlying choice processes (Herrnstein & Prelec, 1991; Soltani & Wang, 2006; Simen & Cohen, 2009), here we show that matching is a robust emergent property of the dopamine system, and that *β* is an intrinsic property of dopamine neurons that can be directly estimated from neuronal recordings. To test whether the model provides the matching law, we model the dynamics of the location of a behaving animal as a stochastic process. Let *R*(*x*) be the input field, which is the expected reward *R* as a function of location *x*. At long time- and length-scales, run-and-tumble motion resembles Brownian motion (Berg, 1993), with a diffusion coefficient *D* ≈ *m*^-1^*ν*^2^*τ*, where *v* is the average run speed, *τ* is the average run duration, and *m* is the dimension (we assume that *d* is close to the adapted level *d*_0_). Brownian motion in the model is biased by longer runs when the agent is moving up the gradient. To account for this, one can add to the diffusion process an advection term that is proportional to the logarithmic gradient: χ∇ log *R*(*x*) (Si *et al*, 2012; Dufour *et al*, 2014; Menolascina *et al*, 2017). Taken together, the stochastic dynamics of the agent are approximated by a Langevin equation similar to the classic Keller-Segel equation used to model chemotaxis (Keller & Segel, 1971):

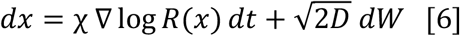

where *W* is an *m*-dimensional Wiener process (see Supplementary Information 1 for the derivation of Eq. 6). For relatively short runs compared with the adaptation time (sub-second order), the advection parameter χ, also called chemotactic drift, is given by: 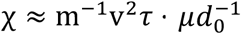(Si *et al*, 2012), and thus rises with velocity v, gain *μ* and mean run duration *τ*. It is important to note that Eq. 6 holds also for the model variant in which dopamine modulates movement speed (see Supplementary Information 2).

Eq. 6 captures how animal movement depends on the parameters of the dopamine circuit, as well as on movement parameters *τ, ν*. Decreasing the average run duration *τ* or average movement speed *ν* (as in Parkinson’s disease) decreases both diffusivity *D* and advection χ, resulting in slower effective motion and gradient climbing. Gradient climbing efficiency (chemotactic drift) increases with 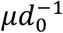, which is the gain of dopamine neurons normalized by their baseline activity. Other circuit parameters do not affect movement dynamics.

Eq. 6 is equivalent to the Langevin Monte Carlo algorithm, a common algorithm from statistical physics for sampling probability distributions (Roberts & Tweedie, 1996; Neal, 2011; Girolami & Calderhead, 2011; Dalalyan, 2014). The steady-state distribution can be readily solved, using standard methods of statistical physics, similar to a Boltzmann distribution in a potential field. The motion samples a probability distribution *P(x)* proportional to a power *β* of the expected-reward distribution:

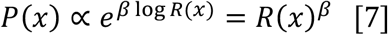

where the power-law *β* equals the normalized gain of the dopaminergic neurons: 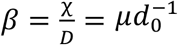. From Eq. 7 we infer that for any two rewards *r*_1_, *r*_2_ the response rates *P*(*x*_1_), *P*(*x*_2_) obey the general matching law presented in Eq. 5.

The reward-taxis model therefore predicts operant matching with 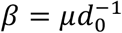. Thus, *β* is equal to the average ratio of gain to baseline activity in dopaminergic neurons. As mentioned above, these values can be estimated from the neuronal measurements of Eshel et al. (Eshel *et al*, 2016), μ ≈ 5 spikes/s and *d*_0_ ≈ 5 spikes/s. These values yield *β* ≈ 1, in agreement with the matching law. Similar parameters are found also in primates (Supplementary Information 3). The agreement is striking since there is no a-priori reason for the gain and baseline to be similar; normalized gain 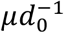 could in principle have a wide range of values including 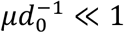 or 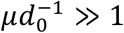.

The matching exponent *β* only depends on parameters that are intrinsic to the dopamine neurons; *β* is independent of movement speed *v* or run duration *τ*, which may vary depending on animal physiology and the environmental context, as well as on the number of dopaminergic neurons. This may explain the robustness of the matching phenomena across species and experimental conditions. Taken together, this suggests that the reward-taxis model may provide a physiological mechanism underpinning operant matching.

### Reward-taxis and pathologies of the dopamine system

Variation and defects in the dopamine system can be classified by their effects on the parameters of the reward-taxis model. Variation in intrinsic neuronal parameters can affect gain μ and baseline firing rate *d*_0_. Such changes are predicted to affect the matching law behavior by changing the exponent *β*. For example, ADHD is associated both with impaired sensitivity to reinforcement (corresponding to low *β*) (Kollins *et al*, 1997; Luman *et al*, 2010) and impaired dopaminergic gain (corresponding to low μ) (Tripp & Wickens, 2008; Hauser *et al*, 2016). Reduced gain can also cause reduced sensitivity to reward gradients so that only strong reward gradients produce continuous focused climbing, as shown by work in bacterial chemotaxis (Long *et al*, 2017). Such hypotheses are in principle testable by detailed behavioral experiments and by experimental manipulation of neuronal gain and baseline properties using drugs or genetic tools.

Other disorders can be due to reduction in the number of functional dopaminergic neurons, as in Parkinson’s disease, where SNc dopaminergic neurons are lost. Such damage is predicted to change mean run duration *τ* and/or mean motion vigor *ν*_0_. If the damage does not sizably affect the intrinsic properties of surviving neurons, such as gain and baseline, they are not expected to change *β* in matching law experiments.

The analogy to chemotaxis can also be used to explore the role of variability in parameters. Variability can occur over time in the same individual (state), or between individuals (trait). In *E. coli* chemotaxis the basal tumbling rate and adaptation time show large non-genetic variability between bacteria that lasts a generation time (Spudich & Koshland, 1976; Levin *et al*, 1998; Levin, 2003; Frankel *et al*, 2014; Keegstra *et al*, 2017; Salek *et al*, 2019). This variation is consequential for gradient climbing performance. Lower basal tumbling frequency improves navigation towards distant targets, whereas high basal tumbling frequency is preferable for nearby targets (Frankel *et al*, 2014). More recently, it was discovered also that pathway gain varies between individual bacteria. Theoretical work indicates that high pathway gain is beneficial for localizing around a single attractant peak, whereas low pathway gain allows efficient crossing between attractant peaks (Karin & Alon, 2021).

Mapping these findings to the dopamine model suggests that state and trait variation can affect reward-taxis. Trait-like variation may be due to developmental or genetic factors. State-like variation in parameters, on the other hand, can potentially be implemented by neuromodulators such as epinephrine and serotonin. Movement vigor shows individuality in humans (Reppert *et al*, 2018), which is in some ways analogous to individuality in baseline tumbling frequency. The reward-taxis model therefore provides testable predictions for the effects of such variations on movement and search behavior.

## Discussion

This study unifies the two main effects of dopamine, encoding reward-derivative and increasing movement vigor, by mapping them to a reward-taxis navigational circuit. The circuit is analogous to the bacterial chemotaxis circuit, where in the dopamine case navigation is along gradients of expected reward. The mapping is based on mathematical analogies at both the physiological and behavioral levels. At the physiological level, experiments on the regulation of dopamine transmission can be parsimoniously explained as fold-change detection (FCD) of expected reward. At the behavioral level, dopamine increases the probability and vigor of movements, thus increasing the duration of correlated motion (“runs”) compared with reorientations (“tumbles”). Both aspects map to the well-characterized chemotaxis navigation circuit in bacteria.

FCD is a recurring motif in biological sensory systems (Adler & Alon, 2018). Therefore, the model of dopamine transmission as FCD of expected reward is in line with the conceptualization of dopamine neurons as sensory neurons for reward (Watabe-Uchida *et al*, 2017). FCD includes the classic Weber’s law of sensory systems, which posits that the maximal response to a change in input is normalized by the background level of the signal (Ekman, 1959). FCD is more general than Weber’s law in that the entire dynamics of the output, including amplitude and duration, is normalized by the background input level. FCD allows the system to function in a scale-invariant manner across several decades of background input (Shoval *et al*, 2010). It also provides a common scale to compare different types of sensory inputs, by referring to their relative (rather than absolute) changes (Hart *et al*, 2013).

While the model focused on the average activity of dopaminergic neurons, the proposed mechanism for FCD (inhibition from neighboring GABAergic neurons) may apply at the level of individual dopaminergic neurons or groups of neurons. This raises the possibility of heterogeneity between dopaminergic neurons – different dopaminergic neurons could become adapted to different expected-reward levels at the same time. This would be consistent with a recent study that demonstrated that a single reward can simultaneously elicit positive and negative prediction errors in different neurons (Dabney *et al*, 2020).

Our model places vertebrate motion regulation in the wider family of run-and-tumble stochastic navigation circuits, which includes motion regulation in bacteria, algae, and simple animals (Pierce-Shimomura *et al*, 1999; Polin *et al*, 2009; Luo *et al*, 2014; Kirkegaard *et al*, 2016). Reward-taxis was anticipated in the early work on TD learning, where Montague et al. showed that run-and-tumble dynamics driven by reward prediction errors can explain observations on bee foraging (Montague *et al*, 1995). Run-and-tumble navigation with FCD guarantees effective sampling of the rewards distributed in the environment, by implementing a search algorithm (Eq. 6) analogous to the widely used Langevin Monte Carlo (LMC) algorithm for sampling probability distributions (Roberts & Tweedie, 1996; Neal, 2011; Girolami & Calderhead, 2011; Dalalyan, 2014) and for global optimization (Chiang *et al*, 1987; Gelfand & Mitter, 1991; Lee *et al*, 2018; Erdogdu *et al*, 2018; Ma *et al*, 2019; Chen *et al*, 2020). The animal samples a probability distribution given by a power of the distribution of expected rewards. This explains the generalized matching law of operant behavior. Furthermore, run-and-tumble navigation provides benefits beyond the Langevin Monte Carlo algorithm by boosting gradient climbing on sufficiently steep reward gradients due to the positive feedback between behavior and sensing. The positive feedback occurs since running along the gradient provides increasing input that further enhances the run duration (Long *et al*, 2017).

The model provides a quantitative and mechanistic explanation of the generalized matching law of operant behavior, with the matching exponent given by the ratio of dopaminergic gain to basal activity. The logarithmic derivative property is crucial for obtaining the matching law. A non-logarithmic derivative, or absolute responses, do not provide matching (Supplementary Information 3).

The model predicts that manipulating the relative gain of dopamine neurons 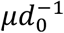 will change the reward sensitivity parameter *β* in the matching law. This prediction can be tested by measuring 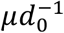 in genetically modified animals where matching behavior is different from wild-type. One such case is mice that are deficient in the cannabinoid type-1 receptor (CB^-/-^), which have *β* that is lower by ∼30% compared with wild-type mice (Sanchis-Segura *et al*, 2004). In agreement with the model prediction, CB^-/-^ mice also have deficient dopaminergic responses relative to baseline (Li *et al*, 2009).

FCD also causes navigation to be invariant (i.e., not affected by) multiplying the input field by a scalar (Shoval *et al*, 2010) – in other words, by multiplying expected rewards across space by a constant factor *λ*. The FCD model predicts that the entire temporal dynamics of the search process, including the distribution of search times, the distribution of run lengths, etc., are invariant to the scale *λ* of reward magnitude. In fact, FCD is a necessary and sufficient condition for scale invariance of a search mechanism (Shoval *et al*, 2010). Scale invariance is prevalent across aspects of perception and behavior (Chater & Brown, 1999; Maylor *et al*, 2001; Stewart *et al*, 2006; Hart *et al*, 2018; Han *et al*, 2020). This property may be crucial for the dopamine circuit, since rewards can vary widely in scale and magnitude.

Together with the similarities there are also differences between bacterial chemotaxis and the reward-taxis model for dopamine. The value of *β* in bacterial chemotaxis is much larger than in the dopamine reward-taxis model, with *β* > 10 in *E. coli* (Hu & Tu, 2014; Karin & Alon, 2021) and *β*∼1 estimated for the dopamine system. The high *β* value in bacteria indicates a strong preference for higher rewards, akin to an optimization for accumulation near attractant peaks. A value of *β*∼1 (which results in the matching law) allows a greater range for exploration of submaximal rewards.

While the presentation of reward-taxis in this study focused on spatial movement, its assumptions are general and it may therefore apply to other behavioral systems. We discussed above the possibility that it applies to hippocampal replay. Another potential system is eye movements. Eye movements are modulated by dopamine and impaired in Parkinson’s disease (Kori *et al*, 1995; Chan *et al*, 2005; Pretegiani & Optican, 2017), and their vigor is modulated by reward prediction errors (Sedaghat-Nejad *et al*, 2019). Since eye movements are commonly studied in behavioral experiments such as reward matching, they may be a good candidate to test the reward-taxis model.

The present minimal model does not include several important aspects including goal-directed behavior and planning. These aspects are likely to complement stochastic navigation with more directed movement. Such a combination of navigation mechanisms is evident in simple organisms that employ run-and-tumble navigation. For example, in *C. elegans* thermotaxis, run-and-tumble navigation is combined with biased reorientations in order to navigate towards an optimal temperature range (Luo *et al*, 2014). Formally, while run-and-tumble navigation resembles Langevin-based sampling, directed reorientations are more closely related to gradient descent, which is efficient for local optimization but poor for global optimization (Ma *et al*, 2019; Raginsky *et al*, 2017; Xu *et al*, 2018). Understanding how animals combine stochastic navigation with other behavioral strategies can therefore improve our understanding of the algorithmic basis of animal behavior.

## Methods

### Model equations and fold-change detection

The equations for dopamine (d) and GABAergic inhibition (g) are provided by:

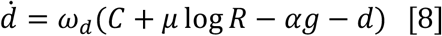

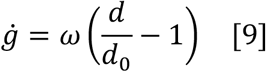

For the dopamine equation, *ω*_*d*_ determines the dopamine degradation rate, *μ*is dopamine gain, *R* is expected reward (defined in the next Methods section), and *α* is GABAergic inhibition strength. For the GABAergic inhibition equation, *ω* determines the adaptation rate and *d*_0_ is the adapted steady-state of dopamine. As is common in FCD circuits, we assume that dopamine dynamics are faster than the dynamics of adaptation due to *g* (i.e., *ω*_*d*_ is large compared with *ω*) so we take:

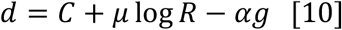

Eqs. 8,9,10 can represent the average activity of individual neurons, or the total activity of many neurons. We therefore used the same equations both to model average individual neuron recordings (as in Figure 1), and to model the effect of dopamine on movement, which is likely to be the sum of the activity of many neurons.

Consider now a constant input *R* = *R*_0_, so that after some time the system reaches steady-state. To find the steady state, we solve Eqs. 8,9, taking 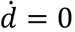, ġ = 0, which yields the steady-state solutions *d*_*st*_ = *d*_0_ and *g*_*st*_ = *α*^−1^(*C* − *d*_0_ + *μ*log *R*_0_). The observation that *d*_*st*_ = *d*_0_ regardless of *R*_0_ and other circuit parameters is an important circuit feature from systems biology known as *exact adaptation* (Barkai & Leibler, 1997; Alon *et al*, 1999; Ma *et al*, 2009; Ferrell Jr, 2016; Alon, 2019). This feature is essential for explaining why dopamine activity returns precisely to baseline after a step increase in expected reward, while GABAergic activity increases in a way that tracks expected reward.

Beyond exact adaptation, the system has an even stronger property of *fold-change detection* (FCD). FCD is defined as dynamics of dopamine (*d*) in response to an input *λR*(*t*) that are independent of *λ*, starting from initial conditions at steady-state for *λR*(0). To show this we replace *g* = *g*′ + *α*^−1^*μ*log *λ*:

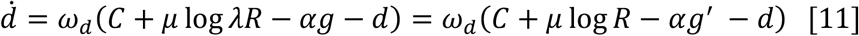

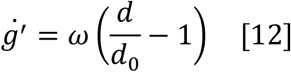

Eqs. 11, 12 are completely independent of *λ*, and their steady-state 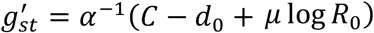 and *d*_*st*_ = *d*_0_ is also independent of *λ*. This means that the dynamics of the system have the FCD property. The FCD property is essential for explaining the scale invariance of the dopaminergic responses to rewards in Fig. 1D – the response only depends on the fold-change of expected reward (two-fold change upon reception of reward at p=0.5) but not on reward magnitude.

While Eqs. 8,9,10 provide FCD, they are not the only possible model that provides FCD for this system. In particular, a feed-forward model where expected reward activates *g* is also possible, i.e.:

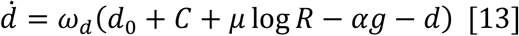

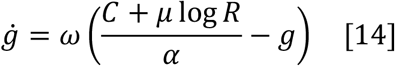

For this circuit, the steady state for a constant input *R* = *R*_0_ is 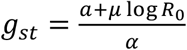 and *d*_*st*_ = *d*_0_. FCD can also be analogously shown. Given an input *λr*(*t*), we can take *g* = *g*′ + *α*^−1^*μ*log *λ*, which again provides equations and steady-state that are independent of *λ*:

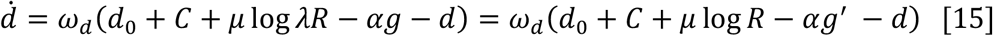

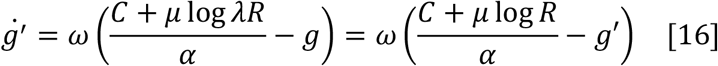

Providing FCD. Parameters used in simulations are provided in Table 1.

**Table 1.**

| Parameter | Value (mouse experiments) | Value (primate experiments) |
| --- | --- | --- |
| $\omega_d$ | 50 s<sup>-1</sup> | 100 s<sup>-1</sup> |
| $\omega$ | 15 s<sup>-1</sup> | 30 s<sup>-1</sup> |
| $\mathcal{C}$ | 15 spike s<sup>-1</sup> | 15 spike s<sup>-1</sup> |
| $\mu$ | 6 spike s<sup>-1</sup> | 6 spike s<sup>-1</sup> |
| $\alpha$ | 0.7 | 0.7 |
| $d_0$ | 5 spike s<sup>-1</sup> | 5 spike s<sup>-1</sup> |

While this simple log-linear model captures various important experimental observations, it is important to note that it has some clear limitations. One limitation is that both *d* and *g* can in principle reach negative values when *R* is small. Measurements of dopamine responses in monkeys indeed show deviations from sub-linearity for small rewards (Stauffer *et al*, 2014). Future studies may build on improved measurements and better mechanistic characterization of the dopamine circuit to refine this model.

### Definition of expected reward and relation between the circuit and TD learning

Here we will define the input to the circuit, which is the logarithmic expected reward log *R*, and present it in the context of the temporal difference (TD) learning theory of dopamine function (Schultz *et al*, 1997; Sutton & Barto, 1998). We first define the expected temporally discounted sum of future rewards *V* (also known as the value function in TD learning):

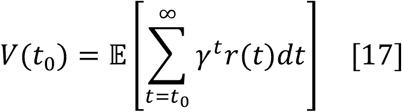

Where *γ* < 1 is a “future discounting” factor and *r*(*t*) is the reward received at time *t* into the future (here for simplicity we take discrete time; for equivalent formulation for continuous time, see Doya et al. (Doya, 2000)). It is possible to think of *V* as a function of the current state *s* of the agent, which may include for example its location in space *x*. This is known as the Markovian setting, where we denote the value function as *V*(*s*). The value function plays an important role in decision making - learning the value function is a principal focus of reinforcement learning algorithms (Schultz *et al*, 1997; Sutton & Barto, 1998). However, for our purposes, we assume that *V* is already learned and can be provided to the circuit as an input. In our model, the input to the circuit for an agent moving into a state *s* at time *t* is defined using the expected reward *R*:

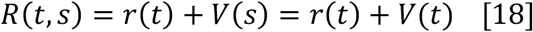

As an example, consider the setting of Figure 1D, where a reward of size *r* = *y* is delivered with probability *p* at Δt time-units into future: the expected reward would in this case be *R*(0) ≈ *pγ*^Δt^*y*. An actual delivery of the reward would then increase *R* to *R*(Δ*t*) ≈ *y*, so the ratio 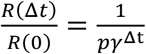 is independent of reward magnitude.

Note that due to discounting and uncertainty, *R* decays with the distance from a location where a reward is delivered, as in Figure 1E (Gershman, 2014).

We will now show that our model is consistent with the TD learning theory of dopamine function. We will first briefly present the TD learning algorithm. In reinforcement learning, the agent usually does not know *V* and needs to learn it from experience. This presents a computational problem, since *V* is an infinite sum over unknown future events. A way to get around this is to update the learned *V* using dynamic programming (Barto *et al*, 1989). The key insight is that Eq. 17 can be rewritten as:

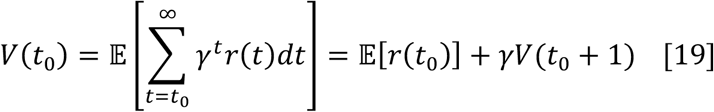

The above equation implies that *V* can be estimated iteratively with a simple update rule, which is at the heart of TD learning. If the agent is at state *s* at time *t*, and at state *s’* at time *t*+*1*, the update rule is:

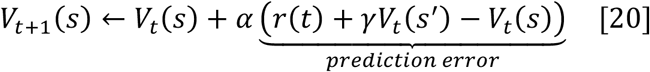

Where *V*_t_(*s*) is the computed estimate of the expected reward at state *s* at time *t, α* is the learning rate, *r*(t) is the reward delivered at time *t* and *γ* is the discounting factor. There is extensive literature demonstrating correspondence between TD learning and midbrain dopamine function (reviewed by (Glimcher, 2011)); specifically, experiments show a correspondence between phasic dopamine secretion and the prediction error term of Eq. 20 (Schultz *et al*, 1997; Glimcher, 2011), in the sense that positive or negative firing of dopamine neurons relative to baseline corresponds positive and negative predictions errors in TD models of learning.

The FCD model proposed here is consistent with the correspondence between dopamine function and TD learning. To see this, notice that learning the logarithm of *V* is equivalent to learning *V*. Also note from Eq. 19:

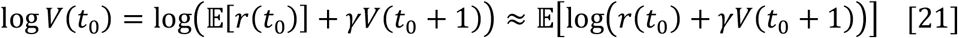

The last approximation holds when *r* is either narrowly distributed, or not very large compared with *γV*(*t*_0_ + 1). Thus, the following update rule is approximately equivalent to the TD update rule, but for log *V*:

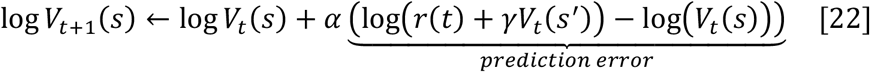

We note that in the continuous limit, and in the absence of reward, the error term is approximately proportional the logarithmic derivative of the value function. This corresponds to the output of our proposed FCD model for low frequency signals.

The output of the circuit to a delivered reward in a transition from a state s to a state s’ is also similar to the above error term:

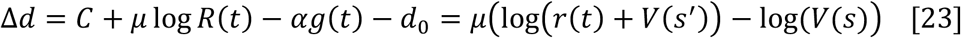

The final equality is due to the fact that prior to reward delivery, GABAergic output adapts to *α*^−1^ (*C* − *d*_0_ + *μ* log *V*(*s*)). The FCD model is therefore compatible with the TD learning theory of dopamine function.

We conclude by noting that since the value that is learned is log *V*, providing the input to the circuit log(*r*(*t*) + *V*(*t*)) requires a rather complex calculation; however, this can be simplified by using the approximation log(*r* + *V*) ≈ max(log *r*, log *V*), which can be implemented by neural circuits (Gawne & Martin, 2002; Knoblich *et al*, 2006).

### Analysis

The fit of the dopaminergic responses in Figure 1B (including confidence intervals) was performed using the NonlinearModelFit function of Mathematica (version 12.1.1). All other figures and simulations were produced using Python (version 3.8.5). The source code and data to produce all the figures is available at https://github.com/omerka-weizmann/reward_taxis.

## Notes

### Competing Interest Statement

The authors have declared no competing interest.

### Summary of Updates

Added code to generate figures and minor corrections.

## References

Adler M & Alon U (2018) Fold-change detection in biological systems. Curr Opin Syst Biol 8: 81–89

Adler M, Mayo A & Alon U (2014) Logarithmic and power law input-output relations in sensory systems with fold-change detection. PLoS Comput Biol 10: e1003781

Adler M, Szekely P, Mayo A & Alon U (2017) Optimal regulatory circuit topologies for fold-change detection. Cell Syst 4: 171–181

Alon U (2019) An introduction to systems biology: design principles of biological circuits CRC press

Alon U, Surette MG, Barkai N & Leibler S (1999) Robustness in bacterial chemotaxis. Nature 397: 168–171

Barkai N & Leibler S (1997) Robustness in simple biochemical networks. Nature 387: 913– 917

Barto AG, Sutton RS & Watkins C (1989) Learning and sequential decision making University of Massachusetts Amherst, MA

Baum WM (1974) On two types of deviation from the matching law: bias and undermatching 1. J Exp Anal Behav 22: 231–242

Baum WM (1981) Optimization and the matching law as accounts of instrumental behavior. J Exp Anal Behav 36: 387–403

Baum WM & Rachlin HC (1969) Choice as time allocation 1. J Exp Anal Behav 12: 861–874

Baum WM, Schwendiman JW & Bell KE (1999) Choice, contingency discrimination, and foraging theory. J Exp Anal Behav 71: 355–373

Beierholm U, Guitart-Masip M, Economides M, Chowdhury R, Düzel E, Dolan R & Dayan P (2013) Dopamine modulates reward-related vigor. Neuropsychopharmacology 38: 1495–1503

Berg HC (1993) Random walks in biology Expanded ed. Princeton, N.J: Princeton University Press

Berg HC & Brown DA (1972) Chemotaxis in Escherichia coli analysed by three-dimensional tracking. Nature 239: 500–504

Berke JD (2018) What does dopamine mean? Nat Neurosci 21: 787–793

Bernoulli D (1968) Specimen theoriae novae de mensura sortis Gregg

Blackwell PG (1997) Random diffusion models for animal movement. Ecol Model 100: 87– 102

Bogacz R (2020) Dopamine role in learning and action inference. Elife 9: e53262

Budick SA & O’Malley DM (2000) Locomotor repertoire of the larval zebrafish: swimming, turning and prey capture. J Exp Biol 203: 2565–2579

Chan F, Armstrong IT, Pari G, Riopelle RJ & Munoz DP (2005) Deficits in saccadic eye-movement control in Parkinson’s disease. Neuropsychologia 43: 784–796

Chater N & Brown GD (1999) Scale-invariance as a unifying psychological principle. Cognition 69: B17–B24

Chen Y, Chen J, Dong J, Peng J & Wang Z (2020) Accelerating Nonconvex Learning via Replica Exchange Langevin Diffusion. ArXiv200701990 Cs Math Stat

Chiang T-S, Hwang C-R & Sheu SJ (1987) Diffusion for Global Optimization in $\mathbb{R}^n $. SIAM J Control Optim 25: 737–753

Codling EA, Plank MJ & Benhamou S (2008) Random walk models in biology. J R Soc Interface 5: 813–834

Cohen JY, Haesler S, Vong L, Lowell BB & Uchida N (2012) Neuron-type-specific signals for reward and punishment in the ventral tegmental area. nature 482: 85–88

Cox J & Witten IB (2019) Striatal circuits for reward learning and decision-making. Nat Rev Neurosci 20: 482–494

Dabney W, Kurth-Nelson Z, Uchida N, Starkweather CK, Hassabis D, Munos R & Botvinick M (2020) A distributional code for value in dopamine-based reinforcement learning. Nature 577: 671–675

Dalalyan AS (2014) Theoretical guarantees for approximate sampling from smooth and log-concave densities. ArXiv Prepr ArXiv14127392

Davidson TJ, Kloosterman F & Wilson MA (2009) Hippocampal replay of extended experience. Neuron 63: 497–507

Davison M (1991) Choice, changeover, and travel: A quantitative model. J Exp Anal Behav 55: 47–61

Davison M & McCarthy D (1988) The matching law: a research review Hillsdale, N.J: L. Erlbaum

Dehaene S (2003) The neural basis of the Weber–Fechner law: a logarithmic mental number line. Trends Cogn Sci 7: 145–147

Dehaene S, Izard V, Spelke E & Pica P (2008) Log or linear? Distinct intuitions of the number scale in Western and Amazonian indigene cultures. Science 320: 1217–1220

Doya K (2000) Reinforcement learning in continuous time and space. Neural Comput 12: 219–245

Dudman JT & Krakauer JW (2016) The basal ganglia: from motor commands to the control of vigor. Curr Opin Neurobiol 37: 158–166

Dufour YS, Fu X, Hernandez-Nunez L & Emonet T (2014) Limits of Feedback Control in Bacterial Chemotaxis. PLOS Comput Biol 10: e1003694

Ek F, Malo M, Åberg Andersson M, Wedding C, Kronborg J, Svensson P, Waters S, Petersson P & Olsson R (2016) Behavioral Analysis of Dopaminergic Activation in Zebrafish and Rats Reveals Similar Phenotypes. ACS Chem Neurosci 7: 633–646

Ekman Gös (1959) Weber’s law and related functions. J Psychol 47: 343–352

Engelhard B, Finkelstein J, Cox J, Fleming W, Jang HJ, Ornelas S, Koay SA, Thiberge SY, Daw ND & Tank DW (2019) Specialized coding of sensory, motor and cognitive variables in VTA dopamine neurons. Nature 570: 509–513

Erdogdu MA, Mackey L & Shamir O (2018) Global Non-convex Optimization with Discretized Diffusions. In Advances in Neural Information Processing Systems 31, Bengio S Wallach H Larochelle H Grauman K Cesa-Bianchi N & Garnett R (eds) pp 9671–9680. Curran Associates, Inc.

Eshel N, Bukwich M, Rao V, Hemmelder V, Tian J & Uchida N (2015) Arithmetic and local circuitry underlying dopamine prediction errors. Nature 525: 243–246

Eshel N, Tian J, Bukwich M & Uchida N (2016) Dopamine neurons share common response function for reward prediction error. Nat Neurosci 19: 479–486

Ferrell Jr JE (2016) Perfect and near-perfect adaptation in cell signaling. Cell Syst 2: 62–67

Frankel NW, Pontius W, Dufour YS, Long J, Hernandez-Nunez L & Emonet T (2014) Adaptability of non-genetic diversity in bacterial chemotaxis. Elife 3: e03526

Friston K (2010) The free-energy principle: a unified brain theory? Nat Rev Neurosci 11: 127– 138

Gawne TJ & Martin JM (2002) Responses of primate visual cortical V4 neurons to simultaneously presented stimuli. J Neurophysiol 88: 1128–1135

Gelfand SB & Mitter SK (1991) Recursive Stochastic Algorithms for Global Optimization in $\mathbb{R}^d $. SIAM J Control Optim 29: 999–1018

Gershman SJ (2014) Dopamine ramps are a consequence of reward prediction errors. Neural Comput 26: 467–471

Girolami M & Calderhead B (2011) Riemann manifold langevin and hamiltonian monte carlo methods. J R Stat Soc Ser B Stat Methodol 73: 123–214

Glimcher PW (2011) Understanding dopamine and reinforcement learning: the dopamine reward prediction error hypothesis. Proc Natl Acad Sci 108: 15647–15654

Gomperts SN, Kloosterman F & Wilson MA (2015) VTA neurons coordinate with the hippocampal reactivation of spatial experience. eLife 4: e05360

Hamid AA, Pettibone JR, Mabrouk OS, Hetrick VL, Schmidt R, Vander Weele CM, Kennedy RT, Aragona BJ & Berke JD (2016) Mesolimbic dopamine signals the value of work. Nat Neurosci 19: 117–126

Han Y, Roig G, Geiger G & Poggio T (2020) Scale and translation-invariance for novel objects in human vision. Sci Rep 10: 1–13

Hanks EM, Hooten MB & Alldredge MW (2015) Continuous-time discrete-space models for animal movement. Ann Appl Stat 9: 145–165

Hart Y, Goldberg H, Striem-Amit E, Mayo AE, Noy L & Alon U (2018) Creative exploration as a scale-invariant search on a meaning landscape. Nat Commun 9: 1–11

Hart Y, Mayo AE, Shoval O & Alon U (2013) Comparing apples and oranges: fold-change detection of multiple simultaneous inputs. PloS One 8: e57455

Hauser TU, Fiore VG, Moutoussis M & Dolan RJ (2016) Computational Psychiatry of ADHD: Neural Gain Impairments across Marrian Levels of Analysis. Trends Neurosci 39: 63– 73

Herrnstein RJ (1961) Relative and absolute strength of response as a function of frequency of reinforcement. J Exp Anal Behav 4: 267

Herrnstein RJ (1970) On the law of effect 1. J Exp Anal Behav 13: 243–266

Herrnstein RJ & Prelec D (1991) Melioration: A theory of distributed choice. J Econ Perspect 5: 137–156

Howe MW, Tierney PL, Sandberg SG, Phillips PE & Graybiel AM (2013) Prolonged dopamine signalling in striatum signals proximity and value of distant rewards. nature 500: 575–579

Hu B & Tu Y (2014) Behaviors and strategies of bacterial navigation in chemical and nonchemical gradients. PLoS Comput Biol 10: e1003672

Karin O & Alon U (2021) Cell-Cell Variation in Chemotaxis Gain Implements a Simulated Tempering Strategy for Efficient Navigation in Complex Environments. Available SSRN 3766494

Keegstra JM, Kamino K, Anquez F, Lazova MD, Emonet T & Shimizu TS (2017) Phenotypic diversity and temporal variability in a bacterial signaling network revealed by single-cell FRET. Elife 6: e27455

Keller EF & Segel LA (1971) Model for chemotaxis. J Theor Biol 30: 225–234

Kim HR, Malik AN, Mikhael JG, Bech P, Tsutsui-Kimura I, Sun F, Zhang Y, Li Y, Watabe-Uchida M, Gershman SJ, et al (2020) A Unified Framework for Dopamine Signals across Timescales. Cell

Kirkegaard JB, Bouillant A, Marron AO, Leptos KC & Goldstein RE (2016) Aerotaxis in the closest relatives of animals. Elife 5: e18109

Knoblich U, Bouvrie J & Poggio T (2006) Biophysical models of neural computation: Max and tuning circuits. In International Workshop on Web Intelligence Meets Brain Informatics pp 164–189. Springer

Kollins SH, Lane SD & Shapiro SK (1997) Experimental analysis of childhood psychopathology: A laboratory matching analysis of the behavior of children diagnosed with attention-deficit hyperactivity disorder (ADHD). Psychol Rec 47: 25– 44

Kori A, Miyashita N, Kato M, Hikosaka O, Usui S & Matsumura M (1995) Eye movements in monkeys with local dopamine depletion in the caudate nucleus. II. Deficits in voluntary saccades. J Neurosci 15: 928–941

Lang M & Sontag E (2016) Scale-invariant systems realize nonlinear differential operators. In 2016 American Control Conference (ACC) pp 6676–6682. IEEE

Lau B & Glimcher PW (2005) Dynamic response-by-response models of matching behavior in rhesus monkeys. J Exp Anal Behav 84: 555–579

Laughlin SB (1989) The role of sensory adaptation in the retina. J Exp Biol 146: 39–62

Lazova MD, Ahmed T, Bellomo D, Stocker R & Shimizu TS (2011) Response rescaling in bacterial chemotaxis. Proc Natl Acad Sci 108: 13870–13875

Lee AK & Wilson MA (2002) Memory of sequential experience in the hippocampus during slow wave sleep. Neuron 36: 1183–1194

Lee H, Risteski A & Ge R (2018) Beyond Log-concavity: Provable Guarantees for Sampling Multi-modal Distributions using Simulated Tempering Langevin Monte Carlo. In Advances in Neural Information Processing Systems 31, Bengio S Wallach H Larochelle H Grauman K Cesa-Bianchi N & Garnett R (eds) pp 7847–7856. Curran Associates, Inc.

Lee RS, Mattar MG, Parker NF, Witten IB & Daw ND (2019) Reward prediction error does not explain movement selectivity in DMS-projecting dopamine neurons. eLife 8: e42992

Levin MD (2003) Noise in gene expression as the source of non-genetic individuality in the chemotactic response of Escherichia coli. FEBS Lett 550: 135–138

Levin MD, Morton-Firth CJ, Abouhamad WN, Bourret RB & Bray D (1998) Origins of Individual Swimming Behavior in Bacteria. Biophys J 74: 175–181

Li X, Hoffman AF, Peng X-Q, Lupica CR, Gardner EL & Xi Z-X (2009) Attenuation of basal and cocaine-enhanced locomotion and nucleus accumbens dopamine in cannabinoid CB1-receptor-knockout mice. Psychopharmacology (Berl) 204: 1–11

Long J, Zucker SW & Emonet T (2017) Feedback between motion and sensation provides nonlinear boost in run-and-tumble navigation. PLoS Comput Biol 13: e1005429

Luman M, Tripp G & Scheres A (2010) Identifying the neurobiology of altered reinforcement sensitivity in ADHD: a review and research agenda. Neurosci Biobehav Rev 34: 744– 754

Luo L, Cook N, Venkatachalam V, Martinez-Velazquez LA, Zhang X, Calvo AC, Hawk J, MacInnis BL, Frank M, Ng JHR, et al (2014) Bidirectional thermotaxis in Caenorhabditis elegans is mediated by distinct sensorimotor strategies driven by the AFD thermosensory neurons. Proc Natl Acad Sci U S A 111: 2776–2781

Ma W, Trusina A, El-Samad H, Lim WA & Tang C (2009) Defining network topologies that can achieve biochemical adaptation. Cell 138: 760–773

Ma Y-A, Chen Y, Jin C, Flammarion N & Jordan MI (2019) Sampling can be faster than optimization. Proc Natl Acad Sci 116: 20881–20885

Maylor EA, Chater N & Brown GD (2001) Scale invariance in the retrieval of retrospective and prospective memories. Psychon Bull Rev 8: 162–167

Mazzoni P, Hristova A & Krakauer JW (2007) Why don’t we move faster? Parkinson’s disease, movement vigor, and implicit motivation. J Neurosci 27: 7105–7116

McDowell JJ (2013) On the theoretical and empirical status of the matching law and matching theory. Psychol Bull 139: 1000

Meder D, Herz DM, Rowe JB, Lehéricy S & Siebner HR (2019) The role of dopamine in the brain-lessons learned from Parkinson’s disease. Neuroimage 190: 79–93

Menolascina F, Rusconi R, Fernandez VI, Smriga S, Aminzare Z, Sontag ED & Stocker R (2017) Logarithmic sensing in Bacillus subtilis aerotaxis. NPJ Syst Biol Appl 3: 16036

Michelot T, Gloaguen P, Blackwell PG & Étienne M-P (2019) The Langevin diffusion as a continuous-time model of animal movement and habitat selection. Methods Ecol Evol 10: 1894–1907

Mohebi A, Pettibone JR, Hamid AA, Wong J-MT, Vinson LT, Patriarchi T, Tian L, Kennedy RT & Berke JD (2019) Dissociable dopamine dynamics for learning and motivation. Nature 570: 65–70

Montague PR, Dayan P, Person C & Sejnowski TJ (1995) Bee foraging in uncertain environments using predictive hebbian learning. Nature 377: 725–728

Morales M & Margolis EB (2017) Ventral tegmental area: cellular heterogeneity, connectivity and behaviour. Nat Rev Neurosci 18: 73–85

Mwaffo V, Anderson RP, Butail S & Porfiri M (2015) A jump persistent turning walker to model zebrafish locomotion. J R Soc Interface 12: 20140884

Neal RM (2011) MCMC using Hamiltonian dynamics. Handb Markov Chain Monte Carlo 2: 2

Nieder A & Dehaene S (2009) Representation of number in the brain. Annu Rev Neurosci 32: 185–208

Nieder A & Miller EK (2003) Coding of cognitive magnitude: Compressed scaling of numerical information in the primate prefrontal cortex. Neuron 37: 149–157

Niv Y, Daw ND, Joel D & Dayan P (2007) Tonic dopamine: opportunity costs and the control of response vigor. Psychopharmacology (Berl) 191: 507–520

Ólafsdóttir HF, Bush D & Barry C (2018) The role of hippocampal replay in memory and planning. Curr Biol 28: R37–R50

Panigrahi B, Martin KA, Li Y, Graves AR, Vollmer A, Olson L, Mensh BD, Karpova AY & Dudman JT (2015) Dopamine is required for the neural representation and control of movement vigor. Cell 162: 1418–1430

Parker NF, Cameron CM, Taliaferro JP, Lee J, Choi JY, Davidson TJ, Daw ND & Witten IB (2016) Reward and choice encoding in terminals of midbrain dopamine neurons depends on striatal target. Nat Neurosci 19: 845–854

Pfeiffer BE & Foster DJ (2013) Hippocampal place-cell sequences depict future paths to remembered goals. Nature 497: 74–79

Pierce-Shimomura JT, Morse TM & Lockery SR (1999) The Fundamental Role of Pirouettes in Caenorhabditis elegans Chemotaxis. J Neurosci 19: 9557–9569

Polin M, Tuval I, Drescher K, Gollub JP & Goldstein RE (2009) Chlamydomonas Swims with Two “Gears” in a Eukaryotic Version of Run-and-Tumble Locomotion. Science 325: 487–490

Pretegiani E & Optican LM (2017) Eye movements in Parkinson’s disease and inherited parkinsonian syndromes. Front Neurol 8: 592

Raginsky M, Rakhlin A & Telgarsky M (2017) Non-convex learning via Stochastic Gradient Langevin Dynamics: a nonasymptotic analysis. ArXiv170203849 Cs Math Stat

Reppert TR, Rigas I, Herzfeld DJ, Sedaghat-Nejad E, Komogortsev O & Shadmehr R (2018) Movement vigor as a traitlike attribute of individuality. J Neurophysiol 120: 741–757

Roberts GO & Tweedie RL (1996) Exponential convergence of Langevin distributions and their discrete approximations. Bernoulli 2: 341–363

Robinson S, Smith DM, Mizumori SJY & Palmiter RD (2004) Firing properties of dopamine neurons in freely moving dopamine-deficient mice: Effects of dopamine receptor activation and anesthesia. Proc Natl Acad Sci 101: 13329–13334

Rubinstein M (1977) The strong case for the generalized logarithmic utility model as the premier model of financial markets. In Financial Dec Making Under Uncertainty pp 11–62. Elsevier

Salek MM, Carrara F, Fernandez V, Guasto JS & Stocker R (2019) Bacterial chemotaxis in a microfluidic T-maze reveals strong phenotypic heterogeneity in chemotactic sensitivity. Nat Commun 10: 1877

Sanchis-Segura C, Cline BH, Marsicano G, Lutz B & Spanagel R (2004) Reduced sensitivity to reward in CB1 knockout mice. Psychopharmacology (Berl) 176: 223–232

Schultz W, Dayan P & Montague PR (1997) A neural substrate of prediction and reward. Science 275: 1593–1599

Sedaghat-Nejad E, Herzfeld DJ & Shadmehr R (2019) Reward prediction error modulates saccade vigor. J Neurosci 39: 5010–5017

Shadmehr R & Ahmed AA (2020) Vigor: neuroeconomics of movement control MIT Press

Shen J (2003) On the foundations of vision modeling: I. Weber’s law and Weberized TV restoration. Phys Nonlinear Phenom 175: 241–251

Shoval O, Goentoro L, Hart Y, Mayo A, Sontag E & Alon U (2010) Fold-change detection and scalar symmetry of sensory input fields. Proc Natl Acad Sci 107: 15995–16000

Si G, Wu T, Ouyang Q & Tu Y (2012) Pathway-Based Mean-Field Model for Escherichia coli Chemotaxis. Phys Rev Lett 109: 048101

da Silva JA, Tecuapetla F, Paixão V & Costa RM (2018) Dopamine neuron activity before action initiation gates and invigorates future movements. Nature 554: 244–248

Simen P & Cohen JD (2009) Explicit melioration by a neural diffusion model. Brain Res 1299: 95–117

Smouse PE, Focardi S, Moorcroft PR, Kie JG, Forester JD & Morales JM (2010) Stochastic modelling of animal movement. Philos Trans R Soc B Biol Sci 365: 2201–2211

Soltani A & Wang X-J (2006) A biophysically based neural model of matching law behavior: melioration by stochastic synapses. J Neurosci 26: 3731–3744

Sourjik V & Wingreen NS (2012) Responding to chemical gradients: bacterial chemotaxis. Curr Opin Cell Biol 24: 262–268

Spudich JL & Koshland DE (1976) Non-genetic individuality: chance in the single cell. Nature 262: 467–471

Stauffer WR, Lak A & Schultz W (2014) Dopamine reward prediction error responses reflect marginal utility. Curr Biol 24: 2491–2500

Steinberg EE, Keiflin R, Boivin JR, Witten IB, Deisseroth K & Janak PH (2013) A Causal Link Between Prediction Errors, Dopamine Neurons and Learning. Nat Neurosci 16: 966– 973

Stella F, Baracskay P, O’Neill J & Csicsvari J (2019) Hippocampal reactivation of random trajectories resembling Brownian diffusion. Neuron 102: 450–461

Stewart N, Chater N & Brown GD (2006) Decision by sampling. Cognit Psychol 53: 1–26

Sutton RS & Barto AG (1998) Introduction to reinforcement learning MIT press Cambridge

Tobler PN, Fiorillo CD & Schultz W (2005) Adaptive Coding of Reward Value by Dopamine Neurons. Science 307: 1642–1645

Tripp G & Wickens JR (2008) Research review: dopamine transfer deficit: a neurobiological theory of altered reinforcement mechanisms in ADHD. J Child Psychol Psychiatry 49: 691–704

Tu Y, Shimizu TS & Berg HC (2008) Modeling the chemotactic response of Escherichia coli to time-varying stimuli. Proc Natl Acad Sci 105: 14855–14860

Watabe-Uchida M, Eshel N & Uchida N (2017) Neural circuitry of reward prediction error. Annu Rev Neurosci 40: 373–394

William BM (1979) Matching, undermatching, and overmatching in studies of choice. J Exp Anal Behav 32: 269–281

Xu P, Chen J, Zou D & Gu Q (2018) Global Convergence of Langevin Dynamics Based Algorithms for Nonconvex Optimization. In Advances in Neural Information Processing Systems 31, Bengio S Wallach H Larochelle H Grauman K Cesa-Bianchi N & Garnett R (eds) pp 3122–3133. Curran Associates, Inc.

Yoon T, Geary RB, Ahmed AA & Shadmehr R (2018) Control of movement vigor and decision making during foraging. Proc Natl Acad Sci 115: E10476–E10485

