## Supplementary Information for "The dopamine circuit as a reward-taxis navigation system"

Karin et al.

**Supplementary Information 1. Derivation of Langevin dynamics and matching law from run-and-tumble model.**

Here we derive the Langevin dynamics for dopaminergic-driven navigation, given by:

$$dx = \chi\,\nabla \log R(x)\,dt + \sqrt{2D}\,dW \quad [1]$$

where $W$ is an $m$-dimensional Wiener process and $R(x)$ is the expected reward field. The advection term is $\chi = \mathrm{m}^{-1}\mathrm{v}^{2}\tau \cdot \mu d_0^{-1}$, where $m$ is the dimension, $v$ is the movement velocity of the animal, $\mu$ is the dopaminergic gain, and $d_0$ is the adapted activity of the dopaminergic neurons. The diffusion term is $D = m^{-1}v^{2}\tau$. The run duration is given by: $\phi = \tau\left(\frac{d}{d_0}\right)$.

The derivation largely follows the derivation of spatio-temporal dynamics of chemotaxis by Si et al. (Si *et al*, 2012), which is provided here for completeness. We will derive here the one-dimensional case (we confirmed the 2D case using numerical simulations). In order to derive the stochastic dynamics of an individual animal, we construct a Fokker-Planck equation for the probability density $P^{\pm}(g,x,t)$ that denotes the probability that an animal with GABAergic activation $g$ is at location $x$ at time $t$, moving at direction $\pm$. We denote the reorientation frequency as $z = \phi^{-1}$. The master equation for $P^{\pm}(g,x,t)$ includes changes due to movement in space, changes in GABA activation, and reorientations:

$$\frac{\partial P^{+}(g,x,t)}{\partial t} = -\frac{\partial (vP^{+})}{\partial x} - \frac{\partial (\dot{g}P^{+})}{\partial g} - \frac{z(g)}{2}(P^{+} - P^{-}) \quad [2]$$

$$\frac{\partial P^{-}(g,x,t)}{\partial t} = \frac{\partial (vP^{-})}{\partial x} - \frac{\partial (\dot{g}P^{-})}{\partial g} + \frac{z(g)}{2}(P^{+} - P^{-}) \quad [3]$$

To simplify the analysis, Si et al. approximate $P^{\pm}$ as a function of the mean methylation level. In the dopamine context, this corresponds to averaging over GABAergic activity $g$, i.e. assuming that all individuals have GABAergic activation level $G^{\pm}(x,t)$ (we then take $\phi$ to be the run duration at that GABA level). Another simplifying assumption is that due to rapid tumbling, the difference between GABA levels $\Delta G(x,t) = \frac{1}{2}\left(G^{+}(x,t) - G^{-}(x,t)\right)$ can be approximated by the difference in average GABA level over the run length:

$$\Delta G \approx -\frac{\partial G}{\partial x} Z^{-1} v \quad [4]$$

Where $G(x,t) = \frac{P_G^{+}G^{+} + P_G^{-}G^{-}}{P_G^{+} + P_G^{-}}$ is the population-weighted average of GABA activation, and $P_G^{+}, P_G^{-}$ correspond to the probability densities integrated over $g$, and we denote $z(G) = Z$.

In order to derive the Fokker-Planck equation, we define density and flux:

$$\rho(x,t) = \int [P^{+} + P^{-}]dg \quad [5]$$

$$J(x) = \int [v(P^+ - P^-)]dg \quad [6]$$

Adding Eqs. 2, 3 and integrating over *g* we get:

$$\frac{\partial \rho}{\partial t} = -\frac{\partial J}{\partial x} \quad [7]$$

Subtracting Eqs. 2, 3, multiplying by *v*, and integrating over *g* we get:

$$\frac{\partial J}{\partial t} = -v^2 \int \frac{\partial}{\partial x}(P^+ + P^-)dg - v\int (\, z(g)P^+ - \, z(g)P^-)dg$$

$$= -v^2 \frac{\partial}{\partial x}\rho - v(\, z(G^+)P_G^+ - \, z(G^-)P_G^-) \quad [8]$$

Linearizing over $G$:

$$z(G^{\pm}) \approx z(G) + \frac{\partial Z}{\partial G}(G^{\pm} - G) \quad [9]$$

We get:

$$z(G^+)P_G^+ - \, z(G^-)P_G^- = z(G)(P_G^+ - P_G^-) + \frac{\partial Z}{\partial G}\left((G^+ - G)P_G^+ - (G^- - G)P_G^-\right)$$

$$= \frac{ZJ}{v} + \frac{\partial Z}{\partial G}\left(\frac{2(\mathrm{G}^+ - G^-)}{\rho}(P_G^+ P_G^-)\right) = \frac{ZJ}{v} + \frac{\partial Z}{\partial G}\left(\frac{\frac{1}{2}(\mathrm{G}^+ - G^-)}{\rho}\left(\rho^2 - \frac{J^2}{v^2}\right)\right)$$

$$= \frac{ZJ}{v} + \frac{\partial Z}{\partial G}\left(\Delta\mathrm{G}\,\rho\left(1 - \frac{J^2}{\rho^2 v^2}\right)\right) \quad [10]$$

We neglect the term $\frac{J^2}{\rho^2 v^2} \ll 1$. Plugging Eq. 10 into Eq. 8, we get:

$$\frac{\partial J}{\partial t} = -v^2 \frac{\partial}{\partial x}\rho - ZJ - v\frac{\partial Z}{\partial G}\Delta\mathrm{G}\,\rho \quad [11]$$

Taking $\frac{\partial J}{\partial t} = 0$, we get that:

$$J = -v^2 Z^{-1}\frac{\partial}{\partial x}\rho - vZ^{-1}\frac{\partial Z}{\partial G}\Delta\mathrm{G}\rho \approx -v^2 Z^{-1}\frac{\partial \rho}{\partial x} - v^2 \frac{\partial Z^{-1}}{\partial G}\frac{\partial G}{\partial x}\rho \quad [12]$$

The final equation is for *G*, which is derived by adding Eqs. 2, 3 multiplying by *g* and integrating over *g*:

$$\int \frac{\partial P^+ g}{\partial t} + \frac{\partial P^- g}{\partial t} dg$$

$$= -\int g\left(\frac{\partial(vP^+)}{\partial x} + \frac{\partial(vP^-)}{\partial x}\right)dg - \int g\left(\frac{\partial(\dot{g}P^+)}{\partial g} + \frac{\partial(\dot{g}P^-)}{\partial g}\right)dg \quad [13]$$

The left-hand side is simply:

$$\int \frac{\partial P^+ g}{\partial t} + \frac{\partial P^- g}{\partial t} dg = \frac{\partial}{\partial t}(P_G^+ G^+ + P_G^- G^-) = \frac{\partial}{\partial t}\rho G = \frac{\partial G}{\partial t}\rho + \frac{\partial \rho}{\partial t} G \quad [14]$$

For the right-hand side:

$$-\int g\left(\frac{\partial(vP^{+})}{\partial x}-\frac{\partial(vP^{-})}{\partial x}\right)dg=-\frac{\partial}{\partial x}v(P_G^{+}G^{+}-P_G^{-}G^{-})=$$

$$=-\frac{\partial}{\partial x}v\big(P_G^{+}G+P_G^{+}(G^{+}-G)-P_G^{-}G-P_G^{-}(G^{-}-G)\big)$$

$$=-\frac{\partial}{\partial x}\left(JG+v\Delta\mathrm{G}\rho\left(1-\frac{J^2}{\rho^2v^2}\right)\right)\approx-\frac{\partial}{\partial x}(JG+v\Delta\mathrm{G}\rho)$$

$$=\frac{\partial\rho}{\partial t}G-\frac{\partial G}{\partial x}J-\frac{\partial}{\partial x}v\Delta\mathrm{G}\rho\ \ [15]$$

And:

$$-\int g\left(\frac{\partial(\dot{g}P^{+})}{\partial g}+\frac{\partial(\dot{g}P^{-})}{\partial g}\right)dg=(\dot{g}(G^{+})P_G^{+}+\dot{g}(G^{-})P_G^{-})\approx\dot{g}\rho\ [16]$$

So we get:

$$\frac{\partial G}{\partial t}\rho+\frac{\partial\rho}{\partial t}G=\dot{g}\rho+\frac{\partial\rho}{\partial t}G-\frac{\partial G}{\partial x}J-\frac{\partial}{\partial x}v\Delta\mathrm{G}\rho$$

$$\frac{\partial G}{\partial t}=\dot{g}-\frac{\partial G}{\partial x}J\rho^{-1}\ -\frac{1}{\rho}\left(\frac{\partial}{\partial x}v\Delta\mathrm{G}\right)\ \ [17]$$

It is assumed that the first term dominates Eq. 17, so dopamine level is close to being adapted $d\approx d_0$ everywhere. This gives $\frac{\partial G}{\partial x}\approx\alpha^{-1}\mu\,\nabla\log R$, and $\frac{\partial\phi}{\partial G}=-\frac{\tau\alpha}{d_0}$, and $Z^{-1}\approx\tau$, so that, by combining Eqs., 7, 12, we get:

$$\frac{\partial\rho}{\partial t}\approx\frac{\partial}{\partial x}\left(v^2Z^{-1}\frac{\partial\rho}{\partial x}+v^2\frac{\partial Z^{-1}}{\partial G}\frac{\partial G}{\partial x}\rho\right)=v^2\tau\left(\frac{\partial}{\partial x}\frac{\partial\rho}{\partial x}-\frac{\partial}{\partial x}\frac{\mu}{d0}\nabla\log R\,\rho\right)\ \ [18]$$

This is the Fokker-Planck equation for the evolution of the probability density of the location of the animal over time, which directly corresponds to Eq. 1 and from which the generalized matching law can be derived.

In the minimal model we considered the case where movement direction was chosen at random after each reorientation. This assumption can be relaxed by allowing for persistence in movement direction after reorientations, or more generally allowing for correlations between the movement direction before and after a reorientation event. Such directional persistence is, in fact, common in run-and-tumble unicellular navigation. While E. coli tumbles result in reorientation angles that are only mildly correlated with the original swimming direction, the reorientations of choanoflagellates only slightly shift their swimming direction (Berg & Brown, 1972; Kirkegaard *et al*, 2016). The effects of directional persistence have been extensively modeled in the literature (Schnitzer, 1993; Locsei, 2007; Kirkegaard & Goldstein, 2018). Roughly, they extend the average run duration $\tau$. Since in our model, changing $\tau$ increases proportionally both the drift term $\chi$ and the diffusion term $\sigma$ in Eq. 1, as long as $\tau$ remains fast compared with the adaptation timescale. This does not affect the conclusions of this work, including the derived matching law, as we verified by numerical simulations.

**Supplementary Information 2. Derivation of Langevin dynamics and matching law from model where dopamine controls movement speed.**

Here we will analyze a model where the reorientation/tumble rate is constant ($z^{-1} = \phi = \tau$) and dopamine controls movement speed: $v(d) = v_0 \frac{d}{d_0}$. We will show that the average dynamics, approximated by Eq. 1, are the same for this model, suggesting that our conclusions generalize to the case where dopamine controls movement speed.

The master equation for this model is:

$$\frac{\partial P^+(g,x,t)}{\partial t} = -\frac{\partial(v(g)P^+)}{\partial x} - \frac{\partial(\dot{g}P^+)}{\partial g} - \frac{\tau^{-1}}{2}(P^+ - P^-) \quad [19]$$

$$\frac{\partial P^-(g,x,t)}{\partial t} = \frac{\partial(v(g)P^-)}{\partial x} - \frac{\partial(\dot{g}P^-)}{\partial g} + \frac{\tau^{-1}}{2}(P^+ - P^-) \quad [20]$$

We again define:

$$\rho(x,t) = \int [P^+ + P^-]dg \quad [21]$$

And:

$$Q(x,t) = \int [P^+ - P^-]dg \quad [22]$$

We will again take the mean field assumption that $P^\pm(g,x,t) = P_G^\pm(x,t)\delta\big(g - G^\pm(x,t)\big)$, with $G^+, G^-$ as the levels of GABA activation of individuals moving in the $+,-$ direction, with the corresponding movement speeds $V^+, V^-$. Adding the equations and integrating over *g*:

$$\frac{\partial \rho}{\partial t} = -\int \left(\frac{\partial(v(g)P^+)}{\partial x} - \frac{\partial(v(g)P^-)}{\partial x}\right) dg = -\frac{\partial}{\partial x}(V^+P_G^+ - V^-P_G^-) \quad [23]$$

Linearizing around the population-weighted average $G$:

$$V^\pm \approx V + \frac{\partial V}{\partial G}(G^\pm - G) \quad [24]$$

We get:

$$\frac{\partial \rho}{\partial t} = -\frac{\partial}{\partial x}\left(VQ + \frac{\partial V}{\partial G}\big((G^+ - G)P_G^+ - (G^- - G)P_G^-\big)\right) = -\frac{\partial}{\partial x}\left(VQ + \frac{\partial V}{\partial G}\Delta \mathrm{G}\rho\left(1 - \frac{Q^2}{\rho^2}\right)\right)$$

$$\approx -\frac{\partial}{\partial x}\left(VQ + \frac{\partial V}{\partial G}\Delta G\rho\right) \quad [25]$$

Subtracting the equations and integrating over *g* provides:

$$\frac{\partial Q}{\partial t} = -\int \left(\frac{\partial(v(g)P^+)}{\partial x} + \frac{\partial(v(g)P^-)}{\partial x}\right) dg - \tau^{-1}Q = -\frac{\partial}{\partial x}(V^+P_G^+ + V^-P_G^-) - \tau^{-1}Q \quad [26]$$

Since we are interested in timescales that go beyond the tumbling rate, we can take the left-hand side of the equation to be zero, so we get:

$$Q = -\tau \frac{\partial}{\partial x}(V^+ P_G^+ + V^- P_G^-) = -\tau \frac{\partial}{\partial x} V\rho \quad [27]$$

Where $V(x,t) = \frac{P_G^+ V^+ + P_G^- G^-}{P_G^+ + P_G^-}$. As before, we will assume that the dynamics are governed by adaptation, so $V \approx v_0$.

Plugging this into Eq. 25 gives:

$$\frac{\partial \rho}{\partial t} = -\frac{\partial}{\partial x}\left(-\tau v_0^2 \frac{\partial \rho}{\partial x} + \frac{\partial V}{\partial G} \Delta G \rho\right) \quad [28]$$

Using the approximation $\Delta G \approx -\frac{\partial G}{\partial x} \tau V$, we get that:

$$\frac{\partial \rho}{\partial t} = -\frac{\partial}{\partial x}\left(-\tau v_0^2 \frac{\partial \rho}{\partial x} - \tau \frac{\partial V}{\partial G} \frac{\partial G}{\partial x} \rho v_0\right) \quad [29]$$

Assuming that dopamine level is close to being adapted $d \approx d_0$, we have $\frac{\partial G}{\partial x} \approx \alpha^{-1} \mu \nabla \log r$, and $\frac{\partial V}{\partial G} = -\frac{v_0 \alpha}{d0}$, so that:

$$\frac{\partial \rho}{\partial t} = v_0^2 \tau \left(\frac{\partial}{\partial x} \frac{\partial \rho}{\partial x} - \frac{\partial}{\partial x} \frac{\mu}{d0} \nabla \log r \, \rho\right) \quad [30]$$

This equation is the same as the dynamics for the case where dopamine controls run duration, suggesting that the average behavior is the same whether dopamine controls movement speed or run duration.

**Supplementary Information 3. Further generalizations of the run-and-tumble model.**

Here we test several other generalizations of the run-and-tumble model, and show that they do not affect our main conclusions. Consider an animal performing run-and-tumble navigation in an environment where the expected reward is $R$, and $d$ is an internal variable that determines run duration $\phi(d)$ and that adapts to a baseline $d_0$. In our case, $d$ is dopamine and adaptation to $d_0$ is due to GABAergic inhibition. The invariant distribution is (Hu & Tu, 2014):

$$P(x) \approx \frac{\Omega}{v} e^{\int_{-\infty}^{x} \left(\frac{\partial \log \phi}{\partial d} \frac{\partial d}{\partial R}\right)_{d=d_0} dR} \quad [31]$$

Where $\Omega$ is a normalization constant, we assume that tumbling is rapid relative to the adaptation of $d$, and we assume that dopamine $d$ is on average close to $d_0$ (see Hu & Tu for derivation details).

Using Eq. 31, we can derive general conditions for which $d$, $\phi$ can provide the general matching law, which requires an invariant distribution such that $\frac{P(x_1)}{P(x_2)} = \left(\frac{r_1}{r_2}\right)^{\beta}$ for rewards of magnitude $r_1, r_2$ at locations $x_1, x_2$, i.e.:

$$\left(\frac{r_1}{r_2}\right)^{\beta} = \frac{P(x_1)}{P(x_2)} = e^{\int_{-\infty}^{x_1} \left(\frac{\partial \log \phi}{\partial d} \frac{\partial d}{\partial R}\right)_{d=d_0} dR - \int_{-\infty}^{x_2} \left(\frac{\partial \log \phi}{\partial d} \frac{\partial d}{\partial R}\right)_{d=d_0} dR} \quad [32]$$

One can see that if dopamine gates the run duration: $\phi \propto \phi_0 d$ then $\left(\frac{\partial \log \phi}{\partial d}\right)_{d=d_0} = \frac{1}{d_0}$, and if dopamine is activated by the logarithm of reward: $d = \mu \log R + C$, then $\frac{\partial d}{\partial R} = \frac{\mu}{R}$ and from Eq. 32:

$$\frac{P(x_1)}{P(x_2)} = e^{\frac{\mu}{d_0} \log r_1/r_2} = \left(\frac{r_1}{r_2}\right)^{\frac{\mu}{d_0}} \quad [33]$$

Providing the matching law with $\beta = \frac{\mu}{d_0}$. It is important to notice that the precise circuit architecture (feedforward or feedback) is not important, only the logarithmic activation of dopamine, and adaptation.

Other possible circuit architectures do not provide the matching law. If dopamine is directly activated by reward $R$, i.e., $d = \mu R^{\alpha} + D$ (with $\alpha > 0$), then $\frac{\partial d}{\partial R} = \alpha \mu R^{\alpha - 1}$ and from Eq. 32:

$$\frac{P(x_1)}{P(x_2)} = e^{\frac{\mu}{d_0}(r_1^{\alpha} - r_2^{\alpha})} \quad [34]$$

Which does not provide the matching law. Another possibility is that dopamine (and hence reorientation frequency) depends on $R$, rather than a gradient of $R$. This possibility, which was analyzed for chemotaxis by Schnitzer (Schnitzer, 1993), results in a uniform (i.e. reward independent) stationary distribution and thus cannot explain the matching law. Finally, movement speed $v$ that depends on the spatial position or on the reward can lead to accumulation near rewards, however to provide a positive relation: $R(x_1) > R(x_2) \rightarrow P(x_1) > P(x_2)$, it is necessary that running speed decreases with dopamine, which is in contrast with the invigorating effect of dopamine on motion.

**Supplementary Information 3.**

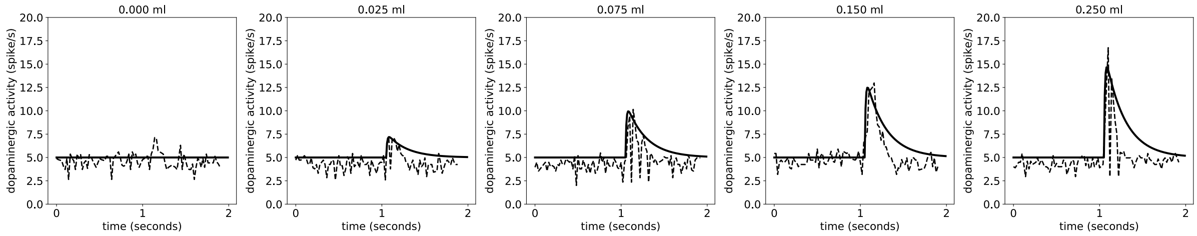


**Supplementary Figure 3. Dopaminergic responses to variable size liquid rewards in monkeys.** Dopamine responses in Macaque monkeys to cues predicting variable size liquid rewards (dashed lines) correspond to model simulations, given by a step $R(t) = R_0 + \lambda u \theta(t - t_0)$ where $\theta(t - t_0)$ is a unit step function, $R_0 = 1$ and $\lambda = 20\ ml^{-1}$, and $u$ is the expected value of the liquid volume that the cue predicts. Data is from the population neuron recordings of Figure 1B in Tobler et al. (Tobler *et al*, 2005).

### References


Berg HC & Brown DA (1972) Chemotaxis in Escherichia coli analysed by three-dimensional tracking. *Nature* 239: 500–504

Hu B & Tu Y (2014) Behaviors and strategies of bacterial navigation in chemical and nonchemical gradients. *PLoS Comput Biol* 10: e1003672

Kirkegaard JB, Bouillant A, Marron AO, Leptos KC & Goldstein RE (2016) Aerotaxis in the closest relatives of animals. *Elife* 5: e18109

Kirkegaard JB & Goldstein RE (2018) The role of tumbling frequency and persistence in optimal run-and-tumble chemotaxis. *IMA J Appl Math* 83: 700–719

Locsei JT (2007) Persistence of direction increases the drift velocity of run and tumble chemotaxis. *J Math Biol* 55: 41–60

Schnitzer MJ (1993) Theory of continuum random walks and application to chemotaxis. *Phys Rev E* 48: 2553

Si G, Wu T, Ouyang Q & Tu Y (2012) Pathway-Based Mean-Field Model for Escherichia coli Chemotaxis. *Phys Rev Lett* 109: 048101

Tobler PN, Fiorillo CD & Schultz W (2005) Adaptive Coding of Reward Value by Dopamine Neurons. *Science* 307: 1642–1645
